# Post-ischemic triiodothyronine treatment improves stroke outcome by stabilizing the blood-brain barrier

**DOI:** 10.1101/2023.11.21.568025

**Authors:** Daniel Ullrich, Dagmar Führer-Sakel, Heike Heuer, Steffen Mayerl, Steffen Haupeltshofer, Linda-Isabell Schmitt, Markus Leo, Tim Hagenacker, Christoph Kleinschnitz, Friederike Langhauser

**Author notes:** **Correspondence** Friederike Langhauser, PhD, Department of Neurology of University Hospital Essen, Hufelandstraße 55, 45147 Essen, Germany.

## Abstract

Thyroid hormones control a variety of processes in the central nervous system and influence its response to different stimuli, such as ischemic stroke. Post-stroke administration of triiodothyronine (T3) has been reported to substantially improve outcomes, but the optimal dosage and time window remain elusive. To this end we investigated the consequences of T3 treatment in an experimental model of ischemic stroke in mice. Our research demonstrated a dose-dependent protective effect of T3 by reducing infarct volumes, with the optimal T3 dosage identified as 25 µg/kg. In addition, we observed a time-dependent effectiveness that was most pronounced when T3 was administered 1 h after transient middle cerebral artery occlusion, with a gradual reduction in efficacy at 4.5 h, and no reduction in infarct volumes when T3 was injected with an 8 h delay. This protective effect persisted for 72 h post-tMCAO, and had accelerated the recovery of motor function by day 3. In-depth investigations further revealed stabilization of the blood-brain barrier, indicated by reduced extravasation of Evans Blue and diminished aquaporin-4 expression, with reduced inflammation and less cell death as underlying reasons. Our findings suggest that thyroid hormones may be a promising intervention for clinical stroke.

## Introduction

Ischemic stroke remains a major global health problem, characterized by its serious consequences and significant societal burden with 12.2 million strokes occurring every year (GBD 2019 Stroke Collaborators). The gold standard treatment for eligible patients is intravenous thrombolysis with tissue plasminogen activator (tPA), administered within the first 4.5 h of symptom onset (Kim et al., 2017). Additionally, mechanical thrombectomy within 24 h after stroke onset is indicated for patients with acute ischemic stroke due to a large artery occlusion in the anterior circulation (Jahdhav et al., 2021). However, these treatment options are only accessible to a very limited number of patients, leading to a high demand for research into new stroke therapies.

There is now increasing evidence that thyroid hormones (TH) play a significant role in the pathophysiology of ischemic stroke. Hypothyroidism has been reported to be a protective factor in ischemic stroke patients (Akhoundi et al., 2011; Alevizaki et al., 2006) and rats (Rastogi et al., 2006), while hyperthyroidism was seen as a risk factor for poor functional outcomes in patients (Wollenweber et al., 2013) and rats (Keshavarz et al., 2016; Rastogi et al., 2008). However, hyperthyroid rats showed profound effects on the cardiovascular system including hypertension and tachyarrhythmia and TH treatment resulted in a catabolic metabolism (Keshavarz et al., 2016), making it difficult to directly link hyperthyroidism to increased infarct volumes.

So far, only a few studies have used TH as a therapeutic treatment in experimental ischemic stroke, indicating a protective effect of THs when administered before (Hiroi et al., 2006; Sadana et al., 2015) or after ischemic stroke (Sadana et al., 2015; Sayre et al., 2017; Genovese et al., 2013). In case of an ischemic insult under euthyroid conditions, acute administration of exogenous TH can avoid neuronal cell damage and reduce infarct volumes, suggesting that timing of modulation of local TH action is crucial. However, these experiments were performed in different species (mice, rats) using different THs (T3, T4, T2, fT3), different TH doses (e.g. 11 µg/kg, 12 μg/kg, 25 µg/kg, 50 µg/kg, 200 µg/kg), times of application (before or after ischemia, single injection or repeated injections), and application routes (intravenous, intraperitoneal).

To this end, we performed a systematic analysis of the effects of THs in ischemic stroke to evaluate the optimal doses and time window for intervention. We subjected male C57BL/6 mice to 60 min of transient middle cerebral artery occlusion (tMCAO), treated them with different doses of T3 at the time of reperfusion and analyzed infarct volumes and neurological deficits. The most effective dose (25 µg/kg) was subsequently injected at different times either before or after stroke induction to evaluate the most effective time window. Mice treated with the most effective dose at the most effective time (1 h after tMCAO) were further analyzed for blood-brain barrier (BBB) integrity, inflammation, thrombosis and cell death.

## Methods

### Study design

This was a randomized, controlled study in 108 male and 20 female C57BL/6N mice (Charles River, Sulzfeld, Germany) at the age of 10-16 weeks. Mice were housed in groups of five in individually ventilated cages at constant room temperature (22°C) and humidity (55±5%), with water and food accessible ad libitum.

Mice were randomly allocated to three groups: sham-operated, vehicle-treated, and triiodothyronine-treated, with group assignment conducted independently. Surgery, behavioral assessments, and evaluation of all readout parameters were performed by an observer blinded to the group allocation, with unblinding subsequent to statistical analysis. Animals meeting any of the following criteria were excluded from the experiment: mortality within 24 h after transient middle cerebral artery occlusion (tMCAO), a Bederson score of 0 at 24 h post-tMCAO, and intracerebral hemorrhage. In total, 110 of the 128 tMCAO-operated animals were included; 18 mice were excluded for one or more of the reasons stated above. The number of animals was calculated via a priori sample size analysis using G*Power software.

All animal experiments were performed in agreement with the ARRIVE (Kilkenny et al., 2010) and IMPROVE guidelines (Percie du Sert et al., 2017), approved by local state authorities (Landesamt für Natur, Umwelt und Verbraucherschutz NRW, LANUV) and conducted in agreement with the German Animal Welfare Act (German Ministry of Agriculture, Health, and Economic Cooperation).

### Transient middle cerebral artery occlusion

Cerebral ischemia was induced through a 60-min tMCAO procedure, as described previously (Langhauser et al., 2012). Mice were anesthetized using 4% isoflurane (Piramal) in 100% O2 for 3 to 5 min (World Precision Instruments, Small Animal Anesthesia System, EZ-7000). Anesthesia was maintained at ∼2% isoflurane, and body temperature was maintained at 37°C using a feedback-controlled warming device (World Precision Instruments, Animal Temperature Controller, ATC-2000). Following a midline skin incision, both the common and external carotid arteries were ligated, while the internal carotid artery (ICA) was temporarily closed using a vascular clip (Fine Science Tools Inc.). Subsequently, a rubber-coated 6.0 nylon monofilament (#6023912, Doccol, Corporation, USA) was inserted into the ICA and advanced into the right internal carotid artery to occlude the origin of the middle cerebral artery (MCA). After 60 min, the filament was carefully withdrawn to achieve recanalization. Animals surviving until day 3 had an occlusion time of 30 min to improve survival. The vascular incision was closed, and the skin incision was sutured. For sham-operated mice, the same surgical procedure was conducted, excluding the insertion of the monofilament.

### Triiodothyronine treatment

In the treatment groups, mice received triiodothyronine (T3) (T6397-100mg; Sigma-Aldrich) at doses of either 10, 25, or 50 µg/kg of body weight, dissolved in 0.9% NaCl solution, either 1 h prior to tMCAO induction or 1, 4.5, or 8 h post-tMCAO induction. In the control group, mice were administered an equivalent volume of 0.9% NaCl. The injections were administered intravenously via the tail vein.

### Calculation of infarct volume and edema

Mice were sacrificed 24 or 72 h after tMCAO and brains were removed and cut into three 2 mm-thick slices (mouse brain slice matrix; Harvard Apparatus). For determination of infarct volume, these slices were immersed in a 2% solution of 2,3,5-triphenyltetrazolium chloride (TTC) in 1× phosphate-buffered solution (PBS) for 10 min at room temperature. Edema-corrected infarct sizes were calculated volumetrically using ImageJ software, provided by the National Institutes of Health, employing the following equation: V_indirect_ (mm^3^) = V_infarct_ × (1 -(V_ih_ - V_ch_) / V_ch_), where (V_ih_ – V_ch_) represents the volume difference between the ischemic hemisphere and the control hemisphere and (V_ih_ – V_ch_)/V_ch_ shows this difference as a percentage of the control hemisphere (Lin et al., 1993).

Brain edema was expressed as a percentage of the normal areas in the contralateral unaffected hemisphere. The extent of swelling was calculated using the Kaplan method: extent of edema = (the volume of right hemisphere – the volume of left hemisphere)/the volume of left hemisphere (Kuts et al., 2019).

### Functional outcomes

Functional assessment was conducted 24 h post-tMCAO using the Bederson score (Bederson et al., 1986) and a modified Neuroscore (Llovera et al., 2015), which is more sensitive than the Bederson score and reflects smaller differences in behavior between treatment groups. The mice were gently placed on a pad and observed for 30 s. Scores were assigned based on the following criteria. For the Bederson score: 0 - No deficits, 1 - Forelimb flexion, 2 - Same as 1 with reduced resistance to lateral pushing, 3 - Unidirectional circling, 4 - Longitudinal spinning, 5 - No spontaneous movements. The modified Neuroscore comprises five key elements, ranges from 0 (no deficits) to 20 (representing the poorest performance in all items), and is calculated as the sum of the general and focal deficits. The Neuroscore results are expressed as a composite neurological score that includes the following general deficits (scores). Spontaneous activity: 0 - The mouse explores its surroundings attentively, 1 - The mouse appears attentive but inactive, 2 - The mouse explores its surroundings intermittently, 3 - The mouse seems dazed and moves sporadically, 4 - No spontaneous movement observed. Body symmetry: 0 - Normal posture, 1 - Slight bending of the body and tail, legs positioned under the body, 2 - Noticeable bending of the body and tail, with legs (partially) extended, 3 - Body and tail bent, mouse lying on its side, 4 - Strongly bent body and tail, mouse lying on its side. Walking: 0 - Smooth gait, 1 - Rigid and slow movement, 2 - Limping or asymmetrical movement, 3 - Shaky and unsteady movement, 4 - No spontaneous movement. Circling: 0 - No circling, 1 - Predominantly unidirectional curves, 2 - Large or predominantly unidirectional circular motion, 3 - Exclusively tight circular motion, 4 - Lateral spinning. Fibril response: 0 - Normal response, 1 - Slow response with contralateral stimulation, 2 - No response with contralateral stimulation, 3 - Same as 2 with slow response to ipsilateral stimulation, 4 - No response bilaterally.

### Fluorescence in situ hybridization

Mice were sacrificed and transcardially perfused with PBS, followed by 4% PFA, 24 hours after tMCAO. Brains were removed and snap-frozen in 2-methylbutane on dry ice. To assess the transition of administered T3 across the blood-brain barrier (BBB) and its activity in the brain, Krueppel-like factor 9 (Klf9) mRNA served as a marker gene for T3 action and was visualized through third-generation fluorescent in-situ hybridization as described elsewhere (Mayerl et al., 2022). Buffers, probes and hairpins were purchased from Molecular Instruments and were applied according to the manufacturer’s instructions. In brief, 20 µm thick coronal brain slices were post-fixed in 4% PFA for 1 hour at 4°C, permeabilized with 0.4% Triton-X100 in PBS for 10 minutes, and dehydrated in ethanol. Sections were incubated in probe hybridization buffer for 10 minutes at 37 °C in a dark and humidified chamber. Probes against KLF9 (commercially available from Molecular Instruments) at a concentration of 0.4 pmol per 100 µl were added to the sections for 24 hours at 37 °C. After rinsing with probe wash buffer and subsequent SSCT (5x SSC + 0.1% Tween-20), sections were incubated with amplification buffer for 30 minutes at room temperature. HCR initiator-matching hairpins h1 and h2 were heat-shocked for 90 seconds at 95 °C and cooled at room temperature in the dark for 30 minutes. Hairpins were added to amplification buffer (6 pmol per 100 µl) and applied onto the slides for 24 hours at room temperature. Slides were washed with SSCT and incubated with 600 nM 4′,6-Diamidin-2-phenylindol (Dapi, D1306, Thermo Scientific) for 10 minutes before final washing in SSCT. Mounting was performed with Fluoromount-G (00-4958-02, Thermo Scientific). Visualization was carried out using Zeiss AxioObserver.Z1 with Zeiss Apotome 3.

### Evans Blue

The tMCAO procedure was conducted as described, with 25 µg/kg T3 or vehicle administered at the time of reperfusion. Two hours prior to sacrifice, 2% Evans Blue (E2129, Sigma-Aldrich) in 0.9% NaCl was intravenously injected into the mice. Following brain removal, the tissue was sliced into 2 mm sections and subsequently post-fixed in 4% paraformaldehyde (PFA) for 30 min at room temperature. These brain slices were then scanned, separated into ipsilateral and contralateral hemispheres, and weighted. The extraction of Evans Blue dye from the tissue was achieved by immersing the whole hemispheres in 500 µl of formamide at 50°C in darkness for a duration of 24 h. Samples were centrifuged for 20 min at 16000 g, and 50 µl of the supernatant was used for fluorometric measurement (excitation wave length 620 nm; emission wave length 680 nm), allowing calculation of the Evans Blue amount per milligram of brain tissue. For each sample, three values were measured and mean values were determined. A calibration line was created by measuring five predefined Evans Blue concentrations.

### Western Blot analyses

Brain slices were divided into ipsilateral and contralateral hemispheres, then further segmented into cortex and basal ganglia. Tissue samples were homogenized in 100 µl of ice-cold RIPA buffer (89900; Thermo Scientific) and treated with ultrasound for 30 s, followed by centrifugation at 16000 g at 4°C for 30 min. The protein concentration was quantified using the biocidin acid assay according to the standard protocol (A55864; Thermo Scientific). Subsequently, 20 µg of each sample was subjected to sodium dodecyl-sulfate-polyacrylamide gel electrophoresis and afterwards transferred onto a nitrocellulose membrane. After a brief 5 min block with Every Blot Blocking Buffer (EBBB; 12010020; Biorad), the membranes were incubated overnight with the following antibodies at 4°C: aquaporin-4 (1:1000; ab81355; Abcam), p-Erk (phosphorylated extracellular signal-regulated kinase)1/2 (1:500; bs-301R; Bioss Antibodies), Erk1/2 (1:1000, 14910882, Invitrogen), and beta-actin (1:10000; ab8227; Abcam). Following thorough washing with TBST (50 mM Tris-HCl, pH 7.4, 0.1% Tween-20), the membranes were incubated at room temperature for 1 h with Alexa Fluor 790-conjugated donkey anti-rabbit (1:5000; A11369; Invitrogen) and Alexa Fluor 647-conjugated donkey anti-mouse antibodies (1:5000; A21240; Invitrogen) in EBBB. Detection was performed using ChemiDoc MP Imaging System (12003154, Bio Rad). For all Western blots, beta-actin was used as a loading control.

### Immunohistology

Mice were sacrificed and transcardially perfused with PBS 24 h after tMCAO and brains were removed and cryo-embedded. To stain CD11b+ and Ly6G+ lymphocytes, cryo-embedded brain sections were fixed in 4% PFA in PBS for 15 min, then blocked for 1 h at room temperature in 5% bovine serum albumin (BSA) with 0.2% Triton-X-100 in PBS, followed by overnight incubation with the primary antibody. The following primary antibodies were diluted 1:100 in PBS with 1% BSA and incubated overnight at 4°C: anti-CD11b (MCA711, Serotec), anti-CD31 (MCA2388, Bio-Rad), anti-GPIX (M051-0, Emfret Analytics), or anti-Ly6G (127601, Biolegend). As secondary antibody, an Alexa 488 conjugated donkey anti-rat antibody (A21208, Life Technologies), dilution 1:100 in PBS with 1% BSA, was used and incubated for 45 min at room temperature. Subsequently, sections were washed in PBS and mounted with a medium containing Mowiol and 1,4-diazabicyclo-(2,2,2)octan to prevent bleaching. For quantification of infiltrated immune cells, five identical brain sections (10 µm) per animal were chosen at the level of the basal ganglia, with a distance of 100 µm, and cells were counted from four different mice from each group. Sections in which the primary antibody had been omitted were used as negative controls. Immune cell-rich spleen sections of mice were used as positive controls.

Apoptotic neurons in the ischemic hemisphere were visualized using TUNEL at 24 h after tMCAO. Brain sections (10 µm thickness) were fixed in acetone for 10 min and blocked for 1 h in 5% BSA in PBS containing 1% goat serum and 0.3% Triton-X100 to prevent nonspecific binding. A mouse antibody to NeuN (MAB377, Millipore, 1:1000 in PBS) was applied overnight at 4°C. Proteins were detected after 45 min incubation in Dylight 488-conjugated goat anti-mouse antibody (ab96871, Abcam) at a dilution of 1:200 in 1% BSA in PBS. TUNEL-positive cells were stained using the in-situ cell death detection kit TMR red (11684795910; Sigma-Aldrich) following the manufacturer’s instructions.

Images were taken with a Leica DMi8 microscope using a Hamamatsu C11440-22 CU camera and the associated Leica Application Suite X Software (LasX 3.0.2.16120). Image processing was performed using ImageJ (National Institutes of Health, USA).

### Statistical analysis

The statistical analysis was performed using GraphPad Prism 8 (GraphPad software). The normal distribution of the data was tested using the D’Agostino and Pearson omnibus normality test. Two groups were compared using the unpaired, two-tailed Student’s t-test. For comparison of multiple groups, 1-way or 2-way analysis of variance (ANOVA) tests with post hoc Bonferroni adjusted t-tests were used for normally distributed data, and Kruskal-Wallis tests with post hoc Dunn multiple comparison tests were used for non-normally distributed data. Non-normally distributed data were presented as scatter plots, with median and normally distributed data as mean ± standard error of the mean (SEM). P values < 0.05 were considered significant (* P < 0.05; ** P < 0.01; *** P < 0.001).

## Results

### Evaluation of most effective T3 dose for intervention 24 h after brain ischemia

C57BL/6 mice underwent a 60 min tMCAO procedure, followed by treatment with different doses of T3 (10 µg/kg, 25 µg/kg, or 50 µg/kg body weight) or 0.9% NaCl (controls) upon reperfusion, aiming to identify the most effective dose for acute-phase intervention. Examination of infarct volumes (TTC staining) revealed a significant reduction in both the 25 µg/kg (mean 38.01 ± 22.64 mm^3^, n = 12; P < 0.001) and 50 µg/kg (mean 40.22 ± 15.88 mm^3^, n = 10; P < 0.01) T3-treated groups 24 h post-tMCAO compared to the controls (mean 76.93 ± 17.10 mm^3^, n = 10). However, the administration of 10 µg/kg T3 failed to yield a statistically significant reduction in infarct volume (mean 54.27 ± 23.31 mm^3^, n = 8; P = 0.081, Fig. 1 A). The reduction in infarct volume correlated with improved functional outcomes, as assessed by the Bederson score, but this did not reach statistical significance (Fig. 1 B). To this end, an additional group of mice was treated with 25 µg/kg T3 vs. NaCl and subjected to the modified Neuroscore evaluation. This revealed significant differences in functional improvement (median score 10 for vehicle-treated (n = 5) vs. 7 for T3-treated (n = 7); P < 0.05) at 24-h post-tMCAO (Suppl. Fig. 1).

**Figure 1:**
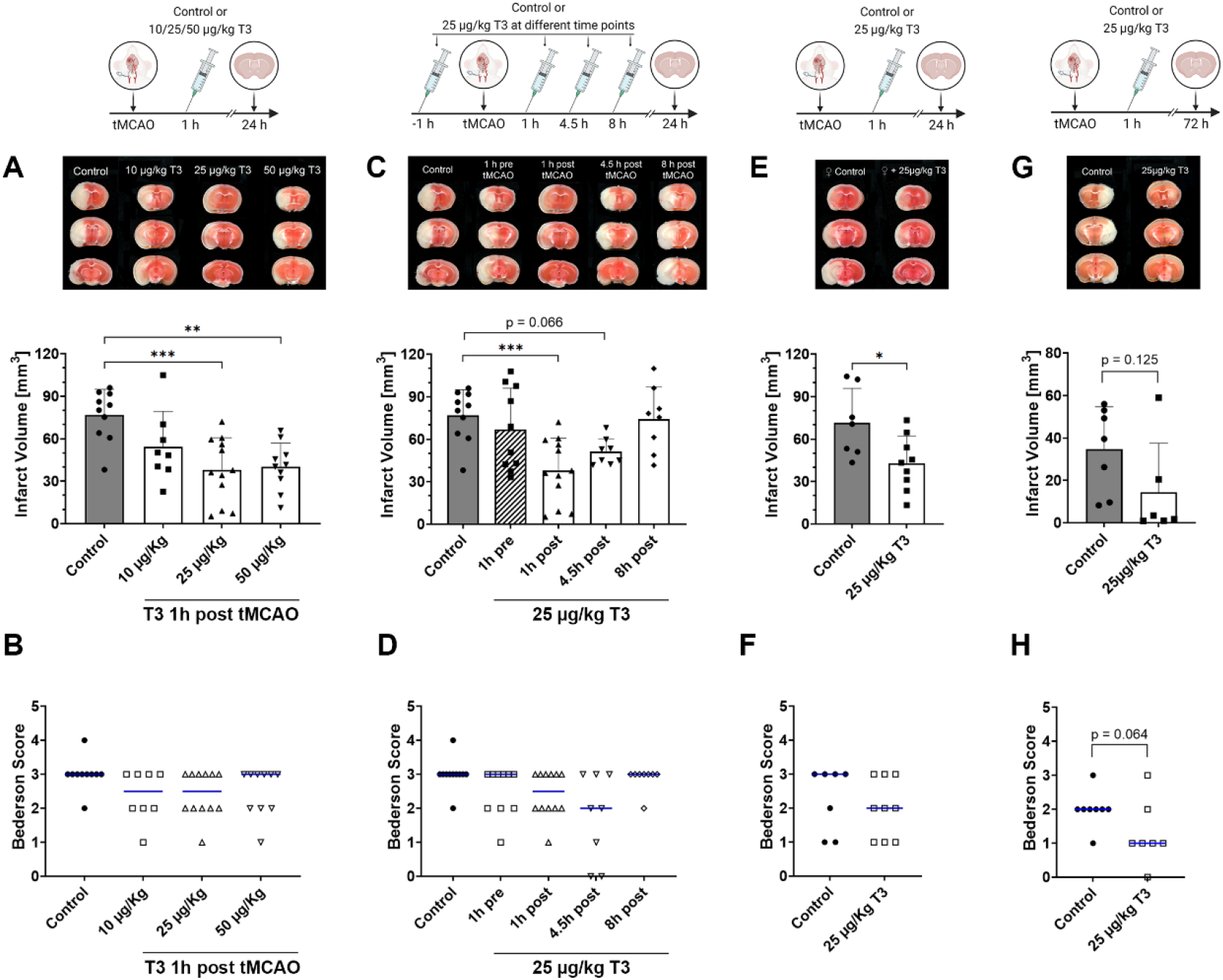
Triiodothyronine (T3) decreases infarct volume after transient middle cerebral artery occlusion (tMCAO) and improves neurological function. **(A)** Representative 2,3,5-triphenyltetrazolium chloride (TTC) stainings of coronal sections from 12-week-old male mice subjected to tMCAO, treated with either 0.9% NaCl as vehicle or 10, 25, or 50 µg/kg T3. Administration of 25 or 50 µg/kg T3 significantly reduced infarct volume (white), while 10 µg/kg had no beneficial effect (n = 7-12 per group;**P < 0.01, ***P < 0.001, 1-way ANOVA). **(B)** The treatment effect was reflected by improved neurological function, indicated by a lower Bederson score. **(C)** Representative TTC stainings of coronal sections of 12-week-old male mice subjected to tMCAO and treated with either 0.9% NaCl as vehicle or 25 µg/kg T3 at different time points (1 h prior to or 1, 4.5, or 8 h after tMCAO). Treatment before tMCAO had no beneficial effect, while administration 1 h after tMCAO had the most significant impact on infarct volume. Administration 4.5 h later still achieved a modest reduction, but after 8 h, no beneficial effect could be observed (n = 7-12 per group; ***P < 0.001, 1-way ANOVA). **(D)** The reduction in infarct volume translated into improved neurological function, as indicated by lower Bederson scores. 12-week-old female mice underwent tMCAO and were subsequently treated with 25 µg/kg T3. TTC staining revealed significantly reduced infarct volumes **(E)** and improved neurological function **(F)** on day 1 (n = 7-9 per group; *P < 0.05, unpaired Student’s t-test). To assess the persistence of the T3 effect, either T3 or vehicle was administered after tMCAO, and the mice were sacrificed on day 3. **(G)** Based on TTC staining, it is evident that T3 still exerted a mild protective effect on day 3 (n = 6-7 per group; P = 0.125, unpaired Student’s t-test), resulting in **(H)** improved neuromotor function.

### Evaluation of the most efficient time window for T3 intervention after brain ischemia

In order to determine the optimal treatment time, we administered the effective dose of T3 (25 µg/kg body weight) at various time points: one hour prior, one hour after, 4.5 hours after, or 8 hours after induction of tMCAO. Administration of T3 before the onset of brain ischemia failed to confer any protective effect (mean 66.83 ± 27.61 mm^3^, n = 10; P = 0.695). The most effective time point for intervention was at the time of reperfusion, 1 h after tMCAO, resulting in a significant reduction in infarct volume (mean 38.01 ± 22.64 mm^3^, n = 12; P < 0.001). Administration of T3 after 4.5 h, the maximal time window for lysis with recombinant tPA (rtPA) in humans, still exhibited a mild reduction in infarct volume (mean 51.01 ± 8.72 mm^3^, n = 8; P = 0.056), albeit less pronounced. However, when administered 8 h after tMCAO, T3 treatment did not yield any alteration in infarct volume (mean 74.11 ± 21.52 mm^3^, n = 8; P = 0.996, Fig. 1 C). The degree of protection correlated with the improvement in neurological outcome, as seen in lower Bederson scores (Fig. 1 D).

To check whether T3 actually reaches the brain, we analyzed the transcription of Krueppel-like factor 9 (Klf9) mRNA as a surrogate marker for T3 transition and activity using fluorescence in situ hybridization. By binding to the cerebrally expressed thyroid hormone receptor alpha, T3 stimulates the transcription of Klf9 mRNA in a concentration-dependent manner. In situ hybridization using probes targeting Klf9 and NeuN revealed a significant 20% reduction in Klf9 transcription in the penumbra on day 1 after tMCAO compared to Sham-operated animals, with no observable change in the contralateral hemisphere. Administration of T3 restored Klf9 transcription in the ipsilateral hemisphere to sham levels and increased it 1.5-fold in the contralateral hemisphere (Suppl. Fig. 2).

### Validation of T3 treatment in female mice

Gender can have a significant impact on stroke outcome in rodents as well as in humans (Roy-O’Reilly and McCullough 2014). To clarify a potential sex difference in T3 effects on stroke size, we also subjected 10- to 16-week-old female C57BL/6 mice to 60 minutes of tMCAO, followed by injection of 25µg/kg T3 or 0,9% NaCl at 1 h after stroke induction. In line with the results in male mice, TTC staining in female mice revealed a notable reduction in infarct volume 24 h post-tMCAO in the T3-treatment group (mean 42.89 ± 19.20 mm^3^, n = 9) compared to the control group (mean 71.43 ± 24.41 mm^3^, n = 7; P < 0.05, Fig. 1 E). This protective effect translated into improved motor function on the first day, as indicated by a reduced Bederson score (Fig. 1 F).

### T3 improved outcome within 72 h after stroke onset

To investigate whether administration of T3 at the time of reperfusion attenuated the damage caused by stroke or merely delayed its occurrence, the outcome after 3 days was examined. In this context, 25 µg/kg of T3 was administered at time of reperfusion, and the mice were monitored daily, until being sacrificed on the third day for infarct volume determination via TTC staining. In contrast to the control group, a reduction in infarct volume following a single administration of T3 (25 µg/kg) was still evident three days after tMCAO, although the reduction was not significant due to one outlier in the T3-treated group (vehicle: mean 34.56 ± 18.61 mm^3^, n = 7; T3-treated group: mean 14.42 ± 21.08 mm^3^, n = 9; P = 0.125, Fig. 1 G). While the T3-treated mice exhibited only mild improvements in motor function on the first day post-tMCAO, the treated animals displayed an enhanced recovery three days later, as evidenced by a reduction in the Bederson score (P = 0.064, Fig. 1 H).

### T3 administration reduced breakdown of the blood-brain barrier

As a next step, we determined whether T3 affects BBB integrity. After ischemia, the blood-brain barrier opens within hours after an infarct, permitting the influx of large and small molecules into the brain. The injection of the azo dye Evans Blue allows to assess the degree of blood-brain barrier breakdown as a consequence. Significantly less Evans Blue was displayed in the brain tissue of T3-treated mice (mean 229.6 ± 42.9 ng/mg, n = 10) compared to controls (mean 306.2 ± 79.1 ng/mg, n = 7; P < 0.05, Fig. 2 A), suggesting stabilization of the BBB in the acute phase.

**Figure 2:**
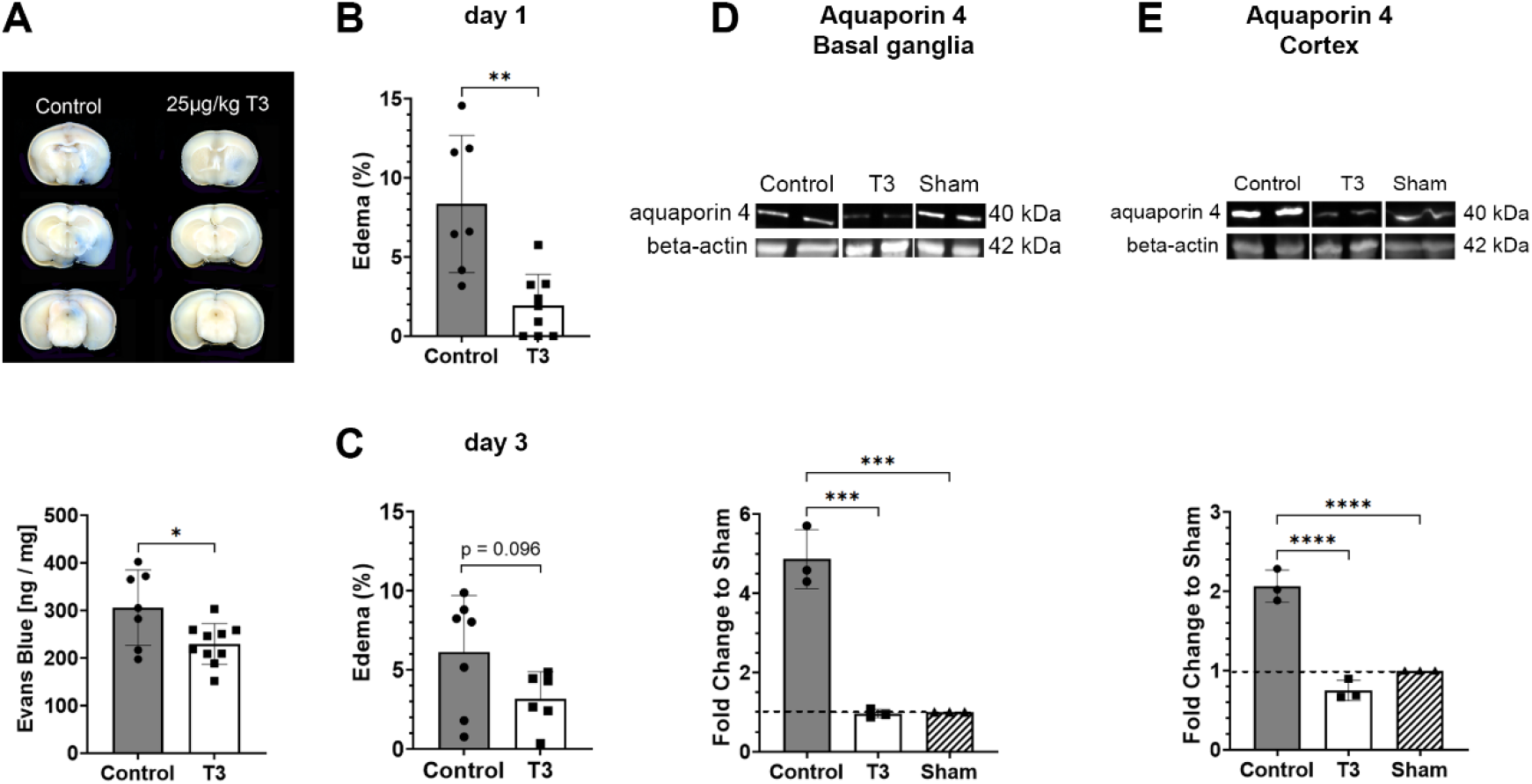
Triiodothyronine (T3) stabilized the blood-brain barrier (BBB) and diminished edema. (**A**) 12-week-old mice underwent transient middle cerebral artery occlusion (tMCAO) and were treated with T3 or 0.9% NaCl as vehicle at the time of reperfusion. Simultaneously, Evans Blue dye was intravenously injected and the mice were sacrificed 24 h later to evaluate the integrity of the BBB. Mice treated with T3 exhibited significantly reduced Evans Blue extravasation compared to vehicle controls (n = 7-10 per group; *P < 0.05, unpaired Student’s t-test). (**B**) Edema was assessed from coronal brain sections. Edema formation on day 1 after tMCAO was significantly diminished following T3 administration compared to controls (n = 7-9; **P < 0.01, unpaired Student’s t-test). (**C**) On day 3, a mild reduction in edema formation was still observed after T3 treatment (n = 6-7 per group; P = 0.096, unpaired Student’s t-test). (**D**) Representative aquaporin-4 Western Blot bands of vehicle, T3-treated, or sham-operated mice after 24 h. Aquaporin-4 expression significantly increased fivefold after tMCAO in the basal ganglia in vehicle-treated mice compared to sham-operated mice. Treatment with 25 µg/kg T3 reduced aquaporin-4 expression to nearly the sham level (n = 3 per group; ***P < 0.001, 1-way ANOVA). (**E**) Aquaporin-4 expression in the cortex doubled in the vehicle controls compared to the sham-operated mice. T3 administration reduced this expression to sham levels (n = 3 per group; ****P < 0.0001, 1-way ANOVA).

In addition, T3-treated mice exhibited significantly less edema after 24 h (mean 1.94 ± 1.84%, n = 9) in contrast to controls (mean 6.46 ± 3.01%, n = 7; P < 0.01, Fig. 2 B). Furthermore, on day 3, the T3-treated mice still demonstrated mildly decreased edema (mean 3.17 ± 1.17%, n = 6) compared to controls (mean 6.10 ± 3.33%, n = 7; P = 0.967, Fig. 2 C), although the reduction was not statistically significant.

To elucidate potential underlying mechanisms of BBB stabilization, aquaporin-4 expression following tMCAO was assessed through Western Blot analysis. Aquaporin-4 expression increased five-fold in the basal ganglia and up to two-fold in the cortex after tMCAO in control mice compared to sham-operated animals. Conversely, on the first day following tMCAO, T3 treatment significantly reduced aquaporin-4 expression, nearly returning to sham levels both in the cortex and basal ganglia (Fig. 2 D + E). This suggests a protective effect of T3 treatment in brain ischemia by stabilizing the BBB.

### T3 administration reduced inflammation, thrombosis, and cell death

To assess the potential impact of T3 treatment on local inflammation and the infiltration of immune cells, immunostainings were conducted 24 hours following stroke onset to visualize Ly6G-positive neutrophil granulocytes and CD11b-positive macrophages/microglia. Following T3 administration, a significant reduction in infiltrated Ly6G-positive neutrophil granulocytes was noted (mean 103 ± 14.4 in controls (n = 3) vs. 28 ± 25.7 in the T3-treated group (n = 4); P < 0.05, Fig. 3 A). However, no discernible differences were observed in the number of CD11b-positive cells (mean 113 ± 15.5 in vehicle controls (n = 3) vs. 93 ± 64.2 in the T3-treated group (n = 4); P = 0.679 (Suppl Fig. 3).

**Figure 3:**
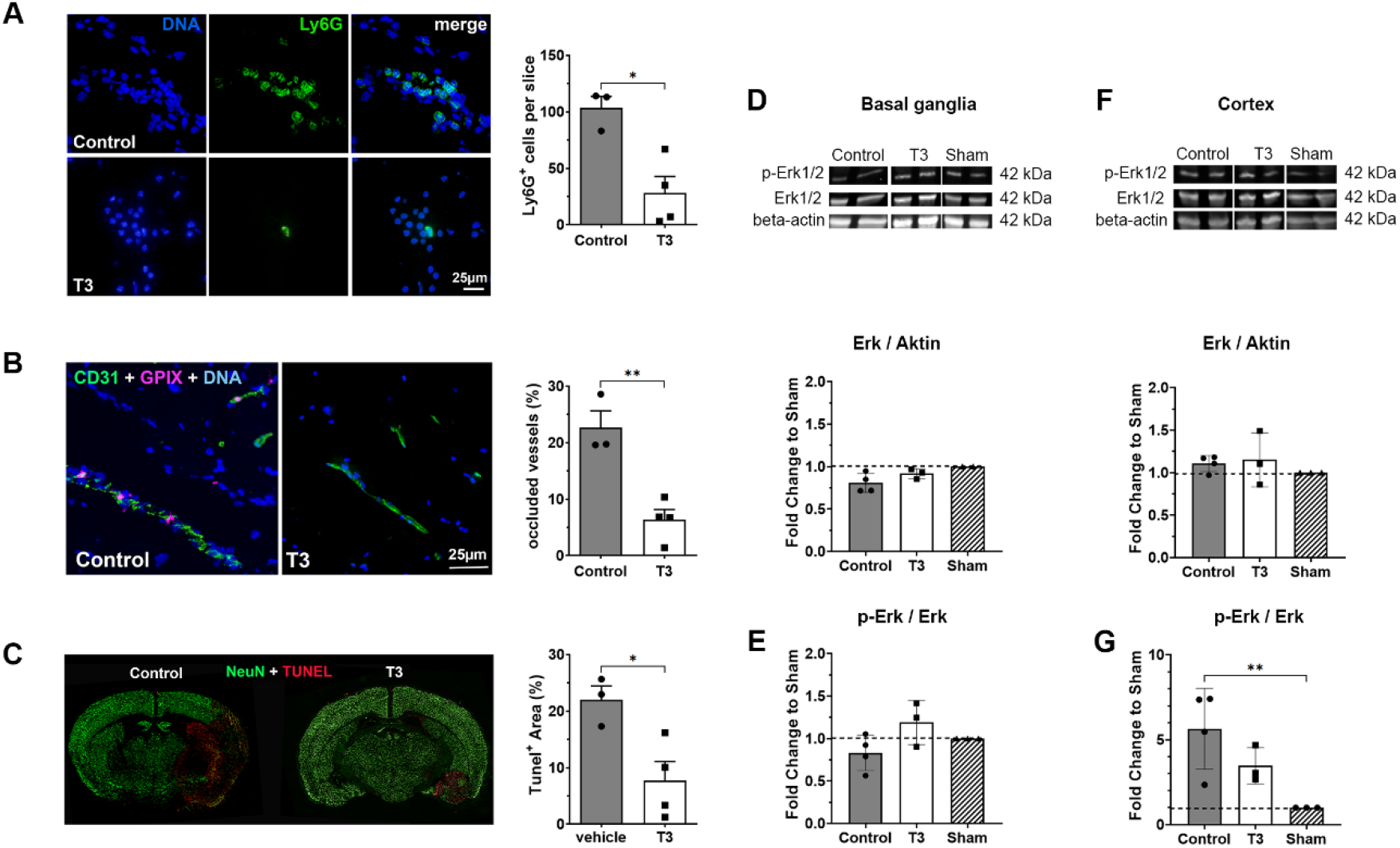
Triiodothyronine (T3) decreases inflammation and apoptosis, and diminishes extracellular signal-regulated kinase (Erk)1/2 activation. Representative fluorescence stainings were conducted to evaluate inflammatory responses and the fraction of occluded microvessels on day 1 after transient middle cerebral artery occlusion (tMCAO). (**A**) The number of Ly6G^+^ neutrophils within the ipsilateral hemisphere was significantly reduced in the T3-treated animals compared to controls (n = 3-4 per group; *P < 0.05, unpaired Student’s t-test). (**B**) The staining for vessels (CD31) and platelets (GP9) revealed that after T3 administration, the proportion of microvessels occluded by thrombi was lower than in vehicle controls. This assessment was conducted in four representative areas within the penumbra (n = 3-4 per group; *P < 0.05, unpaired Student’s t-test). (**C**) To address apoptotic neurons, a cell death assay was conducted together with NeuN immunofluorescence staining. The TUNEL+ area (in red) was quantified and compared between groups. T3-treated mice displayed significantly less apoptosis compared to vehicle controls (n = 3-4 per group; *P < 0.05, unpaired Student’s t-test). (**D + F**) Representative Erk1/2 and phosphorylated Erk1/2 Western blot bands of vehicle, T3-treated, or sham-operated mice on day one after tMCAO were performed. The baseline Erk1/2 expression on day 1 after tMCAO remained consistent in the cortex and basal ganglia across all groups (n = 3-4 per group; P > 0.55, 1-way ANOVA). (**E**) In the basal ganglia, Erk1/2 phosphorylation was unaltered between groups (n = 3-4 per group; P > 0.57, 1-way ANOVA). (**G**) Erk1/2 phosphorylation increased six-fold after tMCAO in the cortex in vehicle-treated mice compared to sham-operated controls. T3 administration reduced the phosphorylation to levels nearly equivalent to those in sham-operated mice (n = 3-4 per group; **P < 0.01, 1-way ANOVA).

Next, we examined whether reduced intracerebral thrombus formation was also one of the underlying mechanisms. In histological brain sections stained for the endothelial marker CD31 and the platelet marker GPIX, about 20% (mean 21.97 ± 4.26%) of all vessels in the ischemic hemisphere were occluded by microthrombi in vehicle-treated mice. In contrast, T3-treated mice showed significantly increased microvascular patency, with less occluded vessels (mean 7.70 ± 6.77% after T3 treatment (n = 4) vs. 22.64 ± 5.14% in vehicle controls (n = 3); P < 0.05, Fig. 3 B).

Furthermore, apoptotic neurons were examined via NeuN and TUNEL double staining to investigate the impact of T3 on DNA fragmentation as a consequence of cell death. The area fraction occupied by apoptotic neurons diminished to one-third after T3 administration (mean % TUNEL+ area 21.97 ± 3.48% in vehicle controls (n = 3) vs. 7.70 ± 3.36% after T3 administration (n = 4); P < 0.05, Fig. 3 C).

As the inhibition of Erk 1/2 is associated with an improved outcome after stroke (C. Schanbacher et al., 2022), the expression and phosphorylation of Erk1/2 were examined in both the cortex and basal ganglia. The expression of Erk1/2 remained constant in all groups following tMCAO (Fig. 3 D + F). In the control group, Erk1/2 phosphorylation significantly increased in the cortex up to six-fold; however, this phosphorylation was strongly attenuated by T3 administration, bringing it to a level that was three times higher than sham expression (see Fig. 3 G). In the basal ganglia, Erk1/2 phosphorylation remained nearly unchanged across all groups after tMCAO (Fig. 3 E).

## Discussion

Our study focused on the potential benefits of T3 administration in a murine model of tMCAO. Treatment with T3 after reperfusion resulted in smaller brain infarcts and enhanced neurological outcomes. We also showed that this effect was independent of sex. Prior research has suggested that administration of TH can improve acute-phase outcomes and decrease infarct volumes after tMCAO (Sadana et al. 2015; Hiroi et al. 2006; Sayre et al. 2016). In our study, we evaluated the dose- and time-dependent effects of T3 application after brain ischemia, which to date has not been fully addressed. While administration of T3 1 h after tMCAO at a dosage of 10 µg/kg did not yield a significant reduction in infarct volume, both 25 and 50 µg/kg showed substantial improvements. The 25 µg/kg dose aligns with that used by Sadana et al. (Sadana et al. 2015), who induced 60 min tMCAO in male CD1 mice and injected 25 µg/kg T3 intravenously 10-15 min after reperfusion. Application of T3 within 4.5 h of tMCAO, which corresponds to the clinical rtPA time window for intervention in humans, still resulted in a mild reduction in infarct volume and improvement in functional outcomes, but had no beneficial effects when given at a later time point. In contrast to other studies (Sadana et al., 2015, Hiroi et al., 2006), we did not observe any protective effect of T3 when administered before stroke onset. This could be due to the fact that the time of pretreatment was 1 h before induction of stroke in our study, but only 30 min in other studies. Another reason could be the use of different mouse strains (C57BL/6 mice in our study and in that by Hiroi et al. vs. CD1 mice used by Sadana et al.) or different times of MCA occlusion (1 h in our study and in that by Sadana et al. vs. 2 h in the study by Hiroi et al.). On the other hand, administration of T3 1 h before tMCAO did not lead to an increase in infarct volume compared to the control animals. Since hyperthyroid rats develop larger infarcts than euthyroid rats (Keshavarz and Dehghani 2017), it could have been that pretreatment with T3 also leads to larger infarcts. However, it appears that a single administration of T3 before stroke onset is not sufficient to worsen stroke outcome. Importantly the benefits of administering T3 1 h after tMCAO persisted for the first three days, showing that T3 treatment did not delay but protected from infarct growth. In addition to the reduction in infarct volume, the treated mice also displayed accelerated recovery of motor function, even though the effect was modest. Whether these protective effects of a single T3 injection can persist over a longer time period remains to be elucidated. So far, there are no studies looking at stroke outcome beyond 24 h, when T3 was administered at the time of reperfusion. There are two studies assessing later times after stroke in the context of TH (Sabbaghziarani et al. 2017; Talhada et al., 2019). These studies focus on regenerative processes such as neuronal regeneration and T3 was administered after the infarct had matured. Consequently, there is no statement on infarct development. A single T3 administration 24 h after tMCAO in rats increased the expression of brain-derived neutrophic factor, Nestin, and SOX2 on day 7 in the subventricular zone (Sabbaghziarani et al. 2017). Talhada and colleagues tested the effects of long-term T3 administration after photothrombosis (Talhada et al., 2019). Intraperitoneal injections of T3 at a dose of 50 µg/kg were started at day 2 post-stroke and were repeated every second day until the end of the experiment on day 14. This treatment significantly improved sensorimotor function in the rotating pole test, without affecting infarct size. Improved motor function was explained by increased dendritic spine density and modulated synaptic neurotransmission by increased levels of synaptotagmin 1&2 and the GluR2 subunit in AMPA receptors and increased dendritic spine density. Enhancement of neurotrophic factors was also seen 24 h after stroke in rats treated with a single dose of thyroxine (Genovese et al., 2013). Taken together, these studies suggest an influence of TH on neuronal regeneration after stroke, and highlight that further pharmacokinetic studies are warranted to fully explore when, how long, or in which interval T3 may exert beneficial effects on ischemic brain injury.

One of the major complications after brain ischemia is the breakdown of the BBB, leading to inflammatory processes, leukocyte infiltration, and the formation of brain edema. Consequently, secondary infarct growth and further deterioration of neurologic symptoms occurs (Ayata and Ropper, 2002; Weiss et al., 2009). Interestingly, the integrity of the BBB was well maintained after T3 treatment, as indicated by reduced Evans Blue extravasation and less edema that persisted over three days. The water channel protein aquaporin-4 is critically involved in the formation of brain edema after ischemic stroke. We identified a reduction in aquaporin-4 expression by T3 in the acute phase of ischemia as one reason for ameliorated edema formation. This is in line with a previous study, where T3 treatment 10-15 min after reperfusion decreased edema formation via a negative regulatory influence on aquaporin-4 expression in CD1 male mice after stroke (Sadana et al. 2015). Negative regulation of aquaporin-4 was also seen in glioblastoma cell lines after stimulation with 50nM T3 (Costa et al., 2019).

Another consequence of a disrupted BBB is an increase in inflammatory processes and leukocyte infiltration. A reduced expression of proinflammatory tumor necrosis factor alpha and interleukin-6 has been observed on day 4 after stroke in rats treated with T3 (Sabbaghziarani et al. 2017). Neutrophils are among the first cells to invade parenchyma and play a causative role in infarct development. Indeed, we observed reduced neutrophil infiltration into the brain parenchyma after T3 treatment, indicating a reduced inflammatory response. Moreover, neutrophils can interact with platelets and endothelial cells and impair tissue reperfusion, a phenomenon commonly referred to as “no reflow phenomenon” (del Zoppo et al., 2003). In our study, a higher microvascular patency with fewer thrombi in the microvasculature was observed in mice treated with T3. An alternative explanation could be a direct effect of T3 on endothelial cells via noncanonical T3 action, as was previously shown by Geist et al. (2022).

Immune cells can also secrete cytokines and stimulate neuronal cell death and promote tissue damage after brain ischemia (Jayaraj et al., 2019). Administration of thyroxine can have anti-apoptotic effects after ischemic stroke in rats by decreasing the expression of pro-apoptotic Bax and increasing anti-apoptotic B-cell lymphoma 2 (Genovese et al. 2013). An anti-apoptotic effect of T3 was also confirmed in our study, as seen by a reduced number of DNA-fragmented cells in the ischemic brain after T3 treatment.

In addition, T3 administration after brain ischemia in rodent models stimulates the phosphoinositid-3-kinase/akt pathway (Hiroi et al. 2006) and suppresses the mitogen-activated protein kinase (MAPK) pathway (Genovese et al. 2013), both of which directly modulate programmed cell death and the inflammatory response. ERK1/2 belongs to the MAPK family and plays a detrimental role in stroke outcome (Schanbacher et al., 2022). ERK is activated by phosphorylation, and this activation was ameliorated by T3 treatment in our stroke model. The inhibition of ERK phosphorylation by T3 was also shown in a model of pressure overload-induced hypertrophied mouse hearts (Suarez et al., 2010).

In summary, in our study we found that a single dose of T3 administered post-stroke reduced infarct volume and improved neurological outcomes, with a lasting effect over 3 days. The rapid onset of beneficial effects strongly suggest non-canonical T3 action. Possible underlying mechanisms of the protective effect include stabilization of the BBB, along with reduced aquaporin-4 expression, a reduced inflammatory response, less cell death, and the suppression of ERK signaling.

## Acknowledgements

This work was supported by the Deutsche Forschungsgemeinschaft (DFG, German Research Foundation) in the framework of SFB/TR 296 LOCOTACT -Project-ID 424957847 (FL, CK, DF, HH and SM) and FOR2789 (CK and FL). The authors thank the Imaging Core Facility Essen (IMCES) for support with the Olympus microscope. We are also grateful to Stefanie Hezel and Kristina Wagner for their dedicated technical support.

## Declaration of Interest

The authors declare that they have no competing financial interests or personal relationships that could have appeared to influence the work reported in this paper.

## Notes

### Competing Interest Statement

The authors have declared no competing interest.

