## Supplemental Figures 1-3 for "Post-ischemic triiodothyronine treatment improves stroke outcome by stabilizing the blood-brain barrier"

Supplementary Figures

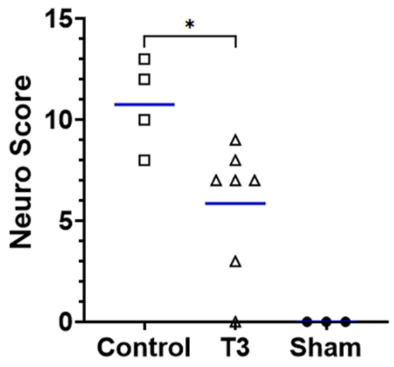


**Suppl Figure 1: Modified Neuroscore.** In order to evaluate the neuromotor function of the mice more precisely, a modified version of the neuroscore was used as a more sensitive behavioural test than the Bederson score. In the five categories assessed, the treated mice consistently showed better neuromotor function than the NaCl-treated controls. The Sham-operated mice displayed no impairments (n = 3-6 per group, *P < 0.05, 1-way Anova).

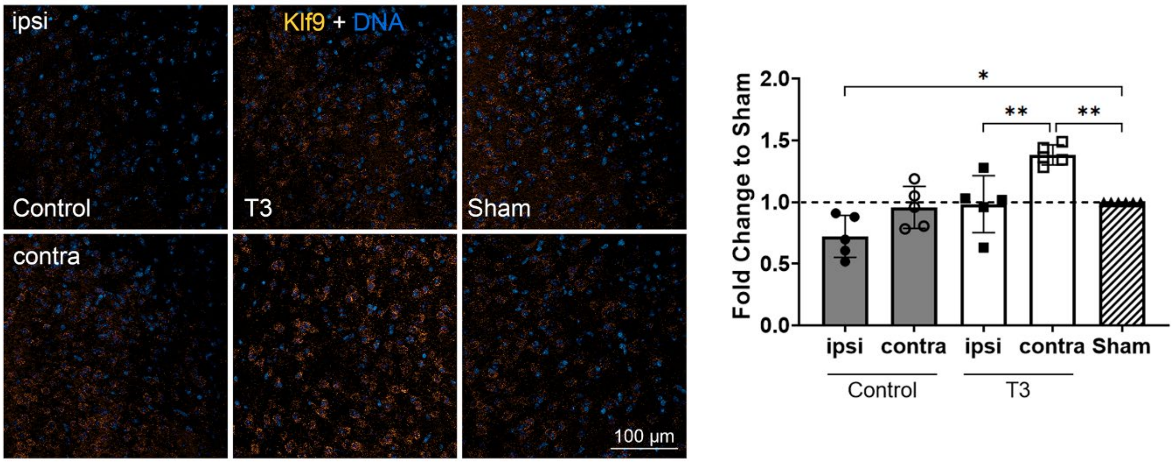


**Suppl Figure 2: Fluorescent in-situ hybridization of Klf9 mRNA as a T3 action marker gene**. Representative third-generation fluorescent in-situ hybridization was performed with probes against Klf9 mRNA as a cerebral T3 reporter to determine crossing of T3 through the BBB and the effect in the brain. Control animals treated with 0.9% NaCl exhibited a significant reduction of Klf9 mRNA transcription in the penumbra on day one after tMCAO compared to sham-operated animals, while the transcription of Klf9 mRNA in the contralateral hemisphere was unchanged. T3 administration restored ipsilateral Klf9 transcription in the penumbra to sham levels on day one. Contralaterally, Klf9 mRNA increased significantly to 1.5-fold sham level in T3 treated mice (n = 5-6 per group, *P < 0.05, **P <0.01, 1-way Anova).

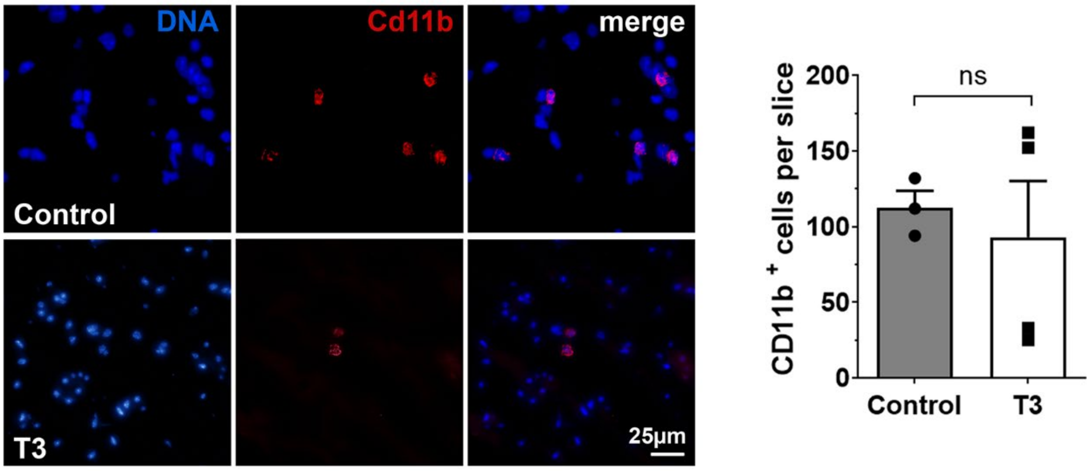


**Suppl Figure 3: Immunofluorescence staining against Cd11b⁺ lymphocytes.** The representative fluorescence staining revealed no difference in the number of Cd11b⁺ lymphocytes in the ipsilateral hemisphere of T3 treated and control mice at day 1 after tMCAO (n = 3-4 per group; *P = 0.679, unpaired Student's t-test).
